# Cue-selective adaptation operates on a separable encoding of stereo and texture information

**DOI:** 10.1101/540260

**Authors:** Evan Cesanek, Fulvio Domini

## Abstract

To perform accurate movements, the sensorimotor system must maintain a delicate calibration of the mapping between visual inputs and motor outputs. Previous work has focused on the mapping between visual inputs and individual locations in egocentric space, but little attention has been paid to the mappings that support interactions with 3D objects. In this study, we investigated sensorimotor adaptation of grasping movements targeting the depth dimension of 3D paraboloid objects. Object depth was specified by separately manipulating binocular disparity (stereo) and texture gradients. At the end of each movement, the fingers closed down on a physical object consistent with one of the two cues, depending on the condition (haptic-for-texture or haptic-for-stereo). Unlike traditional adaptation paradigms, where relevant spatial properties are determined by a single dimension of visual information, this method enabled us to investigate whether adaptation processes can selectively adjust the influence of different sources of visual information depending on their relationship to physical depth. In two experiments, we found short-term changes in grasp performance consistent with a process of cue-selective adaptation: the slope of the grip aperture with respect to a reliable cue (correlated with physical reality) increased, whereas the slope with respect to the unreliable cue (uncorrelated with physical reality) decreased. In contrast, slope changes did not occur during exposure to a set of stimuli where both cues remained correlated with physical reality, but one was rendered with a constant bias of 10 mm; the grip aperture simply became uniformly larger or smaller, as in standard adaptation paradigms. Overall, these experiments support a model of cue-selective adaptation driven by correlations between error signals and input values (*i.e.*, supervised learning), rather than mismatched haptic and visual signals.

## 1. Introduction

### 1.1. Sensorimotor adaptation and 3D shape perception

To ensure that actions produce their intended outcomes, our brains must keep our movements calibrated to the physical environment despite enormous moment-to-moment variability in postural dynamics and neural processing. Over the last 100+ years, research into the process of sensorimotor adaptation has produced a great deal of evidence illuminating when and how the brain adjusts motor planning processes based on sensory feedback (Helmholtz, 1909/1962; Held & Hein, 1958; Welch, 1969; 1978; Cunningham, 1989; Redding & Wallace, 1997; Krakauer, Pine, Ghilardi, & Ghez, 2000; Krakauer, Ghez, & Ghilardi, 2005; Taylor, Krakauer, & Ivry, 2014). However, the role of sensorimotor adaptation in compensating for distortions of 3D shape perception has remained largely unexplored.

When we view and interact with real physical objects, motor control relies on visual information about 3D properties (*e.g.*, slant, curvature, and depth), which are specified by multiple visual cues (*e.g.*, stereo, motion, and texture information). As viewing conditions change from one environment to the next, each of these cues is affected by specific distortions that modulate the amount of bias and noise in their neural encodings. Given the random nature of cue-specific distortions, it is unlikely that the visual system learns to predict the relative accuracies and reliabilities of all depth cues in all possible viewing situations and account for these in the cue-combination process. Thus, rather than providing unwaveringly accurate estimates of 3D shape, visual perception is subject to shifting patterns of inaccuracy as viewing conditions change. For example, an object viewed from a close distance will often be perceived to be deeper than it truly is, due to a constant bias in processing depth from binocular disparities (Johnston, 1991). In another situation, unreliable texture processing could cause significant variable error in the perceived depths of objects with unusual surface markings (Rosenholtz & Malik, 1997; Todd, 2004).

Since the visual guidance of skilled actions like grasping fundamentally depends on estimates of 3D shape, changes in viewing conditions can have real, detrimental consequences on movement accuracy. Consider an observer who suddenly changes the environment where he operates, perhaps moving from a naturally illuminated café during his lunch break to the dimly lit bar where he works. In the café, as he selects a piece of fruit from a nearby bowl, he relies on clear shading gradients and consistent texture patterning on the surfaces of the assorted apples, oranges, and bananas. Later, back in the dim and artificially lit bar, the beer glasses he delivers to patrons are transparent, with unusual specular reflections off their surfaces rather than consistent textures. To avoid fumbling with the glassware, his visuomotor system must quickly recalibrate itself in order to process this new collection of depth cues properly, transforming the available depth information into appropriate grip apertures and orientations when grasping.

The process of sensorimotor adaptation suggests one way to resolve movement errors arising from cue-specific distortions: recalibrate movement planning processes based on sensory feedback from ongoing interactions (Cesanek, Taylor, & Domini, 2019a). However, since multiple depth cues are involved, there is the additional requirement that this recalibration must occur with respect to each individual cue. For example, if stereo information reliably specifies variations in 3D structure, but texture signals are noisy or biased, the relative influence of stereo information in movement planning should be increased. Here, we probed the conditions under which adaptation processes can selectively adjust the influence of individual depth cues when performing actions that require 3D information, following up on promising findings from a previous investigation (Cesanek *et al*., 2019a). We refer to this process as *cue-selective sensorimotor adaptation*, in contrast to the canonical form of adaptation involving a constant shift of motor outputs in one direction.

### 1.2. Identifying the mechanism of cue-selective adaptation

As discussed above, the 3D objects that we grasp and manipulate are always specified by multiple depth cues that can become distorted independently. Thus, the ability to modulate the influence of individual cues from one situation to the next via cue-selective adaptation could be an important capability of biological sensorimotor systems. Experiment 1 was designed to show that the planned grip apertures of grasping movements are affected by cue-selective adaptation when stereo or texture cues to depth suddenly become uncorrelated with physical object depth.

To do this, the sensorimotor system requires a credit assignment mechanism that determines which cue is responsible for detected movement errors. Experiment 2 aimed distinguish between two possible mechanisms for identifying reliable versus unreliable cues and increasing or decreasing their respective influences on the motor output. One possible mechanism involves cross-modal comparisons, where the depth signal from each cue is compared to a haptic “ground truth” signal at the end of each grasp. This mechanism would produce cue-selective adaptation whenever a depth cue fails to indicate the true physical depth. An alternative possibility is an error-based mechanism, which should fail to produce cue-selective adaptation when depth cues are affected by constant biases because these errors can be quickly corrected by canonical sensorimotor adaptation. Under this mechanism, we should observe cue-selective adaptation only when variable errors occur due to an unreliable cue.

The main benefit of the error-based mechanism is that it does not require a ground truth teaching signal to be available in order to reduce the influence of faulty sources of input information. Faulty inputs are identified by their positive correlation with error signals: when a particular cue becomes very noisy, it will take on spurious large values and spurious low values, which will cause positive and negative movement errors, respectively. On the other hand, in the case of a constant bias, there is no systematic variability in the error signals, so it is not possible to identify the faulty cue. As a result, an error-based mechanism would fail to selectively change the influences of the cues. Experiment 2 was designed to test whether cue-selective adaptation occurs (a) even in the case of cue-specific bias or (b) only in response to altered correlations between individual depth cues and physical depth.

In the next section, we describe a model of the process of interacting with 3D objects under cue-specific distortions. This modeling yields two observations that favor the error-based learning mechanism on grounds of ecological function and parsimony. First, we show that canonical adaptation would be sufficient to correct for movement errors when one cue is afflicted by a constant bias, causing the perceived depths of all objects to be uniformly over-or underestimated. In this case, cue-selective adaptation would not be necessary to achieve accurate movements. Second, when canonical adaptation is rendered ineffective by conflicting error signals arising from an unreliable cue, a well-known online supervised learning rule could vary the influence of multiple sources of information based on their long-term correlations with error feedback signals. This mechanism could enable cue-selective adaptation without assuming the availability of directly comparable metric depth estimates from different sensory modalities.

### 1.3. Theoretical model

The visuomotor mapping that mediates grasp planning has previously been shown to be a roughly linear function of object size (Jeannerod, 1981; Säfström & Edin, 2005; but see Smeets & Brenner, 1999 for an alternative view). The output of this function is a target grip aperture, often operationalized as the maximum grip aperture (MGA) of the movement, which occurs just before the hand closes down on the object. At the moment of the MGA, the hand posture must (a) be scaled to the size of the object and (b) provide a “safety margin” so that the fingers enclose the object before clamping down to apply grip force; these two functional characteristics correspond to the slope and intercept parameters of a linear function. To extend this formulation to grasping 3D objects defined by multiple depth cues, we can simply introduce additional slope parameters, which will capture the way that the MGA responds to variations in each of these cues; the intercept remains as a way to introduce a constant offset beyond the observed visual size (*i.e.*, a safety margin).

In this linear model, uniformly shifting the motor output—what we have called canonical adaptation—corresponds to an adjustment of the intercept parameter, whereas cue-selective adaptation is captured by adjustments of the separate slope parameters associated with the available visual cues:

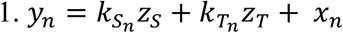

where *y*_*n*_ is the planned motor output on trial *n*, *z*_*S*_ and *z*_*T*_ are the depth values indicated by two distinct visual cues (stereo and texture information), *k*_*S*_*n*__ and *k*_*T*_*n*__ are the slope parameters associated with those cues, and *x*_*n*_ is the intercept parameter. The process of sensorimotor adaptation adjusts the parameters of this mapping according to error feedback signals: canonical adaptation adjusts the intercept, producing constant shifts of the motor output, while cue-selective adaptation adjusts the slopes, altering the relative influences of the available cues on the motor output.

Grasp adaptation is driven primarily by haptic feedback from object contact at the end of each grasp. Specifically, unexpected temporal patterning of the movement, where fingertip contact occurs sooner or later than expected, has been shown to correlate with adaptive changes in the MGA (Säfström & Edin, 2008). Thus, for a given object size, there will be some desired MGA *y*^∗^ that produces the preferred temporal patterning, allowing a stable and comfortable grasp. Errors arise when the actual output does not match this target value. To model this error detection process, we can simply take the difference between the planned motor output *y*_*n*_ and the target output *y*^∗^ preferred for the current physical object depth *z*_φ_, generating a movement error signal ε_*n*_:

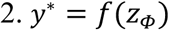

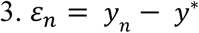

Previous work on reach adaptation has clearly demonstrated that uniform shifts of the intercept parameter are the result of trial-by-trial error corrections, according to some learning rate *b* (Thoroughman & Shadmehr, 2000; Cheng & Sabes, 2006):

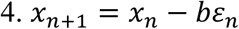

In grasping, this corresponds to a change in the safety margin, applied uniformly across all target objects, regardless of the visual input information. On the other hand, changes of the cue-specific slope parameters *k*_*S*_*n*__ and *k*_*T*_*n*__, which scale the motor output in accordance with the visual inputs, are not often considered in the context of sensorimotor adaptation. To gain some leverage on this question, we can begin by asking when such adjustments would be necessary to support accurate motor performance.

Applying the error-correction process described by Equations 2–4 to the standard reach adaptation experiment described in section 1.2, we find that rightward errors (*y*_*n*_ > *y*^∗^) sensibly cause future movements to aim more to the left, while leftward errors (*y*_*n*_ < *y*^∗^) induce shifts to the right. We demonstrated previously that this same process is sufficient to correct errors that occur due to simple over-or underestimation of depth from a particular cue (Cesanek *et al*., 2019). This is because the faulty cue introduces a relatively constant bias to all motor outputs, pushing them away from the desired output in one direction. Therefore, constant biases do not demand cue-selective adaptation. Cue-selective slope changes become necessary for accurate motor performance only when the strength of the correlation between a given cue and physical reality is reduced, since the unreliable cue will sometimes bias outputs toward depth overestimation and sometimes toward underestimation. The remedy for this more complex pattern of error signals is to reduce the relative influence of this faulty cue.

To see how the necessary slope changes could arise through an error-based learning mechanism, consider the pattern of error signals that would occur across different values of an unreliable cue: spurious high values will misleadingly increase the motor output, causing positive errors, whereas spurious low values will decrease the motor output, causing negative errors. Thus, error signals will be positively correlated with the values of an unreliable cue. This fact can be exploited to perform slope adjustments with a well-known rule for online supervised learning:

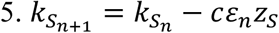

where *C* is a learning rate, *z*_*S*_ is the input from a given depth cue (in this case, stereo), and *k*_*S*_*n*__ is the associated slope parameter on trial *n*. Notice that the product in the second term will be, on average, positive for an unreliable cue (prior to the subtraction). This learning rule works most efficiently if error signals are centered on zero, creating a sensible complementarity with the rapid intercept adjustments that compensate for constant errors. Critically, this error-based learning mechanism for cue-selective adaptation predicts that interfering error signals are necessary to elicit slope changes (*e.g.*, the grip is too large on some trials, but too small on other trials). In contrast, this mechanism would not be expected to operate when one cue is distorted by a constant bias. A biased cue would result in similar movement error signals for all target objects—in this case, the mechanism described by Equation 5 would produce proportional decreases in the influence of each cue, which are indistinguishable from the intercept shifts of Equation 4.

As an alternative hypothesis, it is also possible that the sensorimotor system compares metric depth estimates derived from each cue with a similar metric estimate derived from haptic feedback when the object is stably held. By doing so, the system could identify the biased cue and subsequently reduce its influence in the visuomotor mapping. Indeed, cross-modal comparisons have been tentatively suggested as a possible mechanism of adjusting relative cue influences (Ernst, Banks, & Bülthoff, 2000). Under this hypothesis, we would expect to observe cue-selective adaptation when participants grasp objects defined by one accurate cue and one biased cue.

### 1.4. Study overview

In a tabletop virtual reality environment (with consistent accommodative and vergence information), participants repeatedly grasped 3D objects (paraboloids) defined by stereo and texture cues. At the end of each grasp, the hand closed down on a real object with a physical depth that was set to match the depth specified by one or both of the cues, depending on the feedback condition. Our results indicate that cue-selective adaptation reliably occurs when a single depth cue suddenly becomes uncorrelated with physical reality (and is therefore positively correlated with movement errors). On the other hand, when the depth specified by a particular cue is biased so as to under-or overestimate the physical depth, cue-selective adaptation is absent, but motor outputs are shifted uniformly toward the reinforced cue.

## 2. Results

### 2.1. Experiment 1

Figure 1a depicts a three-by-three matrix of 3D paraboloid objects, where rows of the matrix correspond to different values of texture-specified depth and columns correspond to the different values of stereo-specified depth. Along the main diagonal, we obtain three *cue-consistent* stimuli, where the two cues are rendered based on the same physical depth value. The six off-diagonal stimuli are *cue conflicts*: texture depth is greater than stereo depth in the lower-left region, whereas stereo depth is greater than texture depth in the upper-right. In Experiment 1, we recruited 25 participants to repeatedly grasp these nine stimuli along their depth dimension in two conditions: a haptic-for-texture condition and a haptic-for-stereo condition, where the depth specified by the indicated cue always matched the physical object encountered at the end of the grasp. Consequently, the other cue was uncorrelated with physical depth.

**Figure 1.**
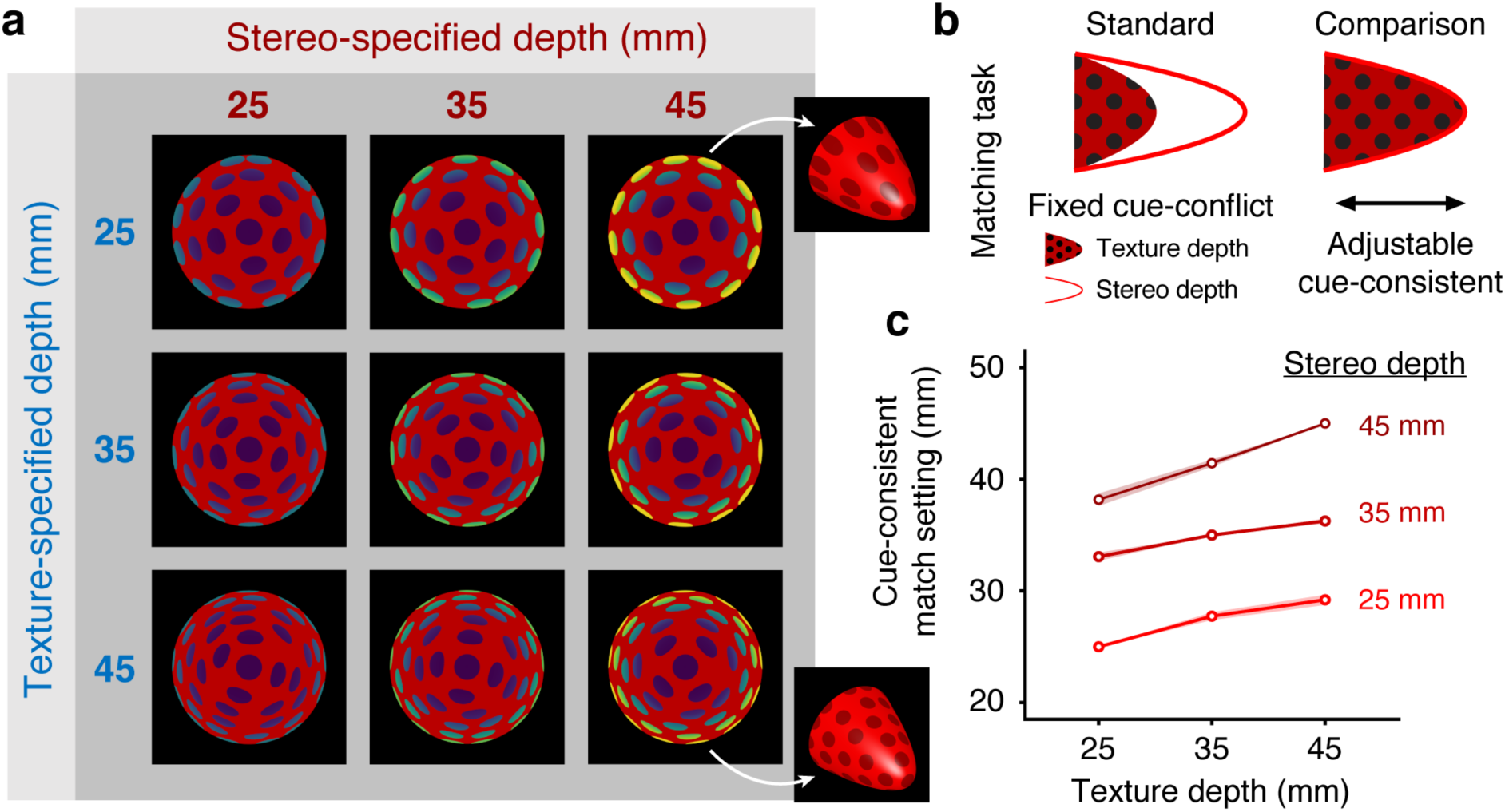
Experiment 1: Stereo-texture paraboloid stimuli and Matching task results. (A) Nine paraboloid objects were rendered by independently manipulating texture and stereo cues to specify depths of 25, 35, or 45 mm (base always subtended 8°). For ease of viewing, stereo depth is coded by a color gradient. The main diagonal of the matrix corresponds to the normally occurring covariation of stereo and texture information (*i.e.*, consistent cues). In this set of stimuli, the effect of variations in stereo information can be assessed independently of the effect of variations in texture via 2D linear regression (Eqn. 1). The off-diagonal objects are generated with cue conflicts. Two oblique views of the final rendered 3D objects are shown on the far right—the dots are circular on the cue-consistent stimulus (bottom-right), while the dots appear stretched on the cue-conflict stimulus (top-right) such that the frontally viewed projection of the texture specifies a shallower stimulus. (B) At the beginning of each session, participants performed a depth adjustment task to obtain a cue-consistent perceived depth match with each of the cue-conflict stimuli (polka dot pattern indicates texture depth, red dotted line indicates stereo depth). In this task, participants alternated freely between viewing one of the conflict objects (the standard, fixed during each trial) and viewing/adjusting the current setting of an adjustable cue-consistent object (the comparison). For each participant, this task identified a set of cue-consistent stimuli that were perceived to have the same depths as the cue-conflicts. In the Grasping task, these cue-consistent stimuli were presented in a Baseline phase to calibrate grasping behavior prior to introducing the cue conflicts, and afterwards in a Washout phase. (C) Average depth setting of the cue-consistent object when adjusted to match the perceived depth of each stereo-texture conflict object. The cue-consistent stimuli are plotted as reference points. Errors ribbons are ±1 SEM.

To obtain a set of cue-consistent stimuli that could be used to calibrate grasping behavior before introducing the cue conflicts, we asked participants to perform a Matching task at the start of each session (Figure 1b). For each of the six cue-conflicts from the test set (note that Matching was not necessary for the three cue-consistent stimuli), they adjusted the depth of a cue-consistent paraboloid until the two appeared to have the same depth. Each participant grasped objects from their personalized set of perceptually matched cue-consistent stimuli in a Baseline phase before the test stimuli were introduced, and again afterwards in a Washout phase. The average depth settings from the Matching task are shown in Figure 1c; these settings correspond to an average relative weight on stereo information *w*_*S*_ of 0.75 (SEM = 0.019) according to *w*_*S*_ = (*z*_*match*_ − *z*_*T*_)/(*z*_*S*_ − *z*_*T*_). Notice that here we have used cue weights that sum to one, instead of freely varying coefficients as in our grasp planning model. This is because our psychophysical procedure relied on a comparison with a fixed standard, and thus cannot indicate the exact metric depth that was perceived—an independent metric probe would be required to do this. The Matching procedure only allows a measurement of the relative influences of the two cues in perception.

We adopted the Matching procedure primarily because it allowed us to obtain a precise perceptual match for each of the target objects in a relatively small number of trials. Having obtained these, we were then able to test in the Grasping task whether the perceived depth matches were treated as such by the visuomotor system, or if a switch from cue-consistent to cue-conflict stimuli would cause an immediate change in grasp performance. Figure 2 plots the MGAs for the Baseline phase (cue-consistent stimuli) against those for the first nine trials (*i.e.*, first bin) of the Adaptation phase (cue-conflicts). The fact that the values are nearly identical across the switch suggests that the input to the visuomotor system was, by and large, the same as the perceptual encoding used for the Matching task. The cue-consistent depths presented during Baseline account for 97% of the variability in Baseline grip apertures, compared to 95% of the variability in early Adaptation (adjusted R^2^), suggesting that grasp planning was based on highly similar encodings of depth information in both phases.

**Figure 2.**
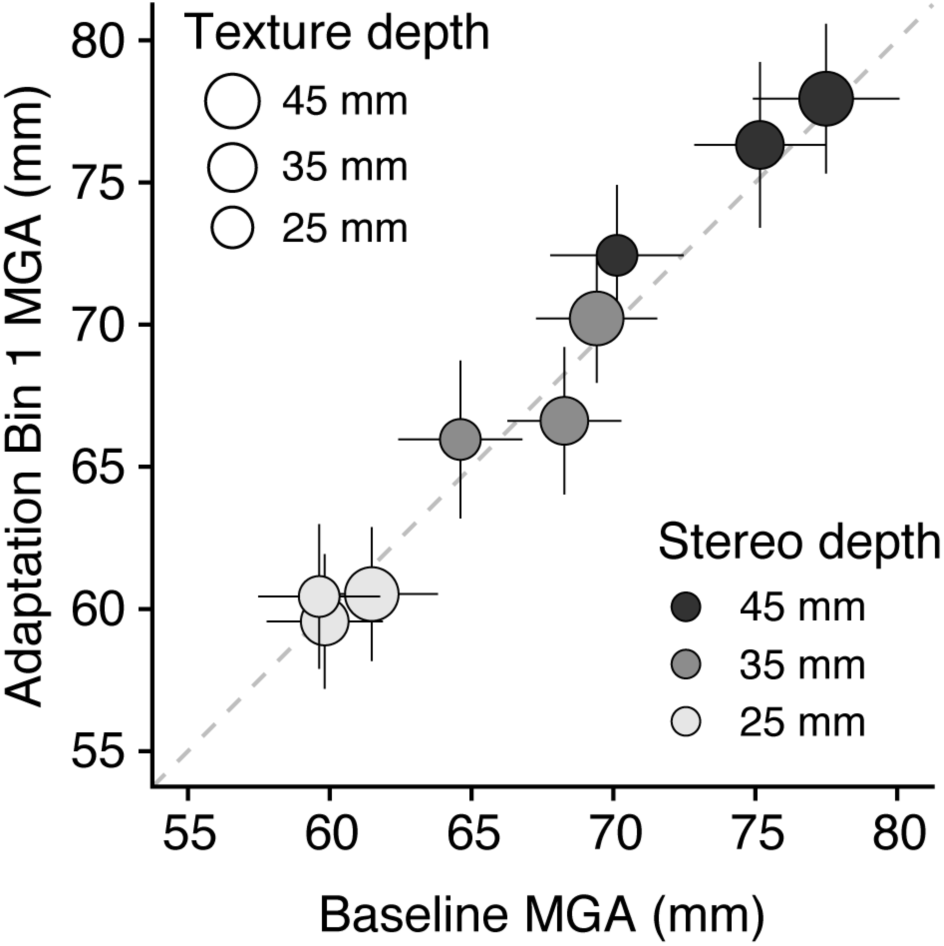
Comparison of maximum grip apertures (MGAs) during Baseline and in the first bin of the Adaptation phase. Across this transition, the component stereo and texture depths of each stimulus changed from consistent to conflicting, but the perceived depth of each object remained the same due to the Matching procedure (see Fig. 1 for average cue-consistent depths). The strong correlation between the MGAs supports the idea that the visuomotor system relies on the same analysis of depth as the perceptual Matching task.

Most importantly, cue-selective adaptation was revealed by changes in the slope of the MGA with respect to stereo-specified and texture-specified depth over the course of Adaptation (Fig. 3). A three-way repeated-measures ANOVA (Condition ✕ Bin ✕ Cue) revealed a significant main effect of Cue (*F*(1, 24) = 60.98; p < 0.0001), representing the stronger influence of stereo information, and a significant three-way interaction (*F*(1, 24) = 5.36; p = 0.029). The latter statistic is the critical one with respect to cue-selective adaptation: it reflects our finding that the difference between stereo and texture slopes becomes smaller over time in the haptic-for-texture condition, and larger over time in the haptic-for-stereo condition. A follow-up two-way ANOVA restricted to the texture slopes yielded no significant effects (all p > 0.5), whereas restricting the test to the stereo slopes yielded a significant interaction of Condition ✕ Bin (*F*(1,24) = 5.14; p = 0.033). This indicates that the three-way interaction of the omnibus test was driven primarily by opposing changes in the stereo slope, with no noticeable changes in the texture slope. This finding was supported by a bin-by-bin analysis of the coefficients, shown in Figures 3a and 3d. When analyzing the difference in these estimates of stereo slope change per bin across haptic feedback conditions, we found that the rate of change was significantly modulated by condition (one-tailed *t*-test; *t*(24) = 2.20; p = 0.019). Linear regressions on the stereo slopes as a function of bin number estimated an average change of +0.01 per bin in the haptic-for-stereo condition, and an average change of −0.01 per bin in the haptic-for-texture condition.

**Figure 3.**
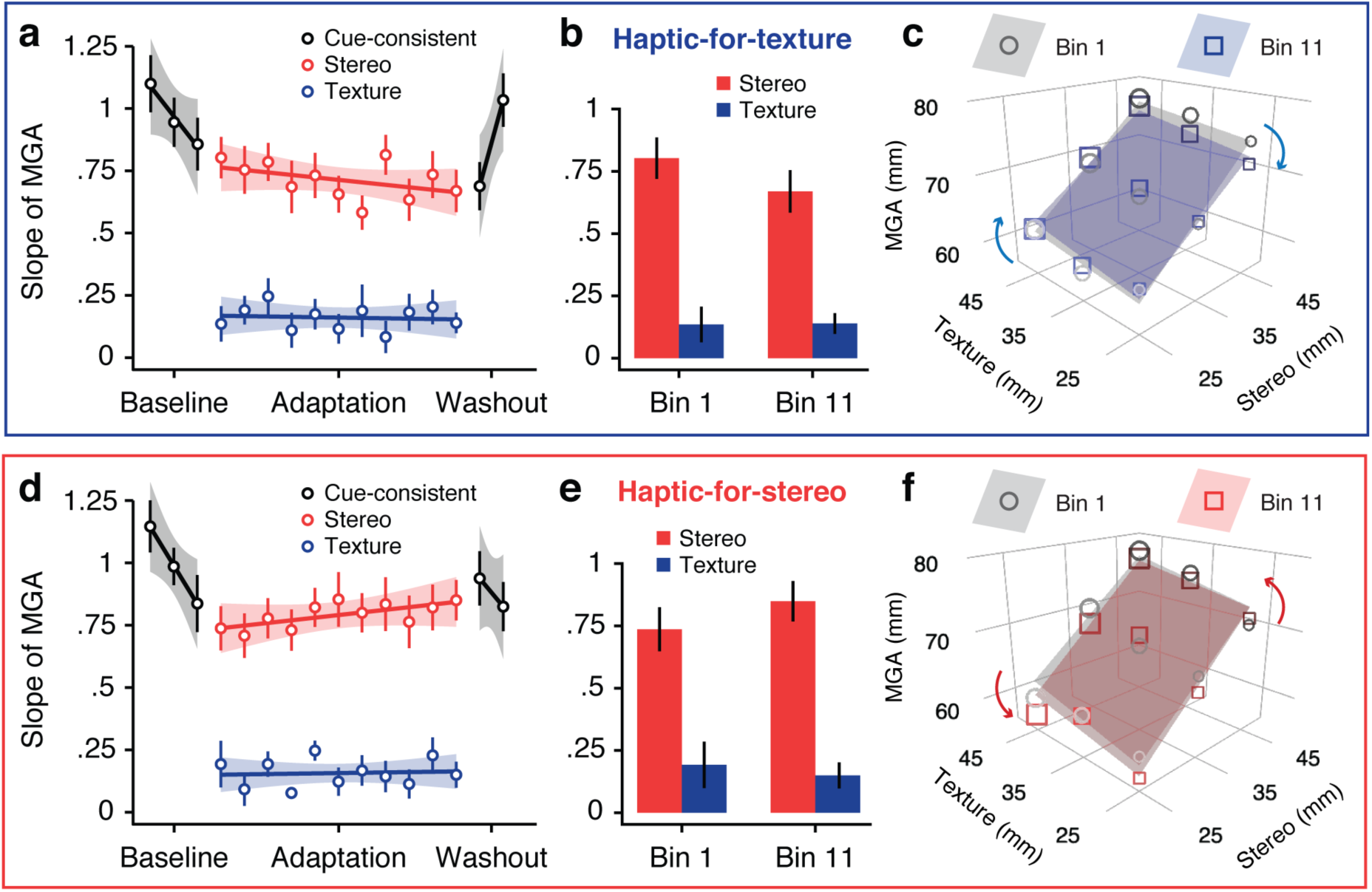
Cue-selective adaptation in Experiment 1. Top panel: Haptic-for-texture condition. Bottom panel: Haptic-for-stereo condition. (a, d) Slope parameters estimated by linear regression on maximum grip apertures (MGAs) as a function of depth information in each bin. For the cue-consistent stimuli presented during Baseline and Washout, we computed a single slope in a simple linear regression. For the cue-conflicts presented during Adaptation, we computed independent slopes with respect to the rendered stereo and texture depths in a multiple regression (Eqn. 1). To evaluate whether the influence of stereo and texture information changed in response to the haptic feedback within the Adaptation phase, we fit a further linear regression on these estimated slopes as a function of bin number. (b, e) Stereo and texture slopes in the first and last bins of Adaptation. (c, f) Surface plots of the 2D regression results during Baseline (gray) and Adaptation (blue or red), plotted over the average MGA for each of the nine target objects. In the haptic-for-texture condition, MGAs shifted down in regions of the stimulus space where stereo misleadingly specified more depth than texture, and shifted up in regions where stereo misleadingly specified less depth than texture. The haptic-for-stereo condition elicited opposite shifts.

Our bin-by-bin slope analysis also indicates a small aftereffect on the slope of the MGA with respect to the cue-consistent depths presented in Baseline and Washout, where the changes in the stereo coefficient that occurred during Adaptation appear to have carried over into Washout, also reducing or increasing the slope of the MGA across the cue-consistent stimuli. In the haptic-for-texture condition, the first bin of Washout showed a smaller slope than Baseline, whereas in the haptic-for-stereo condition, the first bin of Washout showed a slope similar to Baseline. We found a trend toward a slope aftereffect, measured as the effect of feedback condition on the change in cue-consistent slope from Baseline to the first bin of Washout (*t*(24) = 1.45; p = 0.079). Additionally, the slopes in the first bin of Washout differed significantly across condition (*t*(24) = 2.00; p = 0.028). Since aftereffects are typically considered a hallmark of sensorimotor adaptation, this trend provides converging evidence for our main claim of an adaptive change in the relative influence of stereo and texture information within the visuomotor mapping.

### 2.2. Experiment 2

As discussed in the Introduction, cue-selective adaptation can be modeled as a change in the slope parameters of the visuomotor mapping, whereas standard adaptation paradigms have focused on shifts of the intercept. Intercept adjustments are sufficient for eliminating movement errors that arise when depth from one or more cues is over-or underestimated. In contrast, slope adjustments are necessary only when a cue changes the strength of its correlation with physical reality, as this results in movement errors that tend to be positive for spuriously large values of the noisy cue and negative for spuriously low values. Faced with conflicting error signals across the domain of visual inputs, uniform shifts of the intercept would oscillate unhelpfully, but slope adjustments can produce increases in some regions of the visual input space and decreases in other regions, as seen in Experiment 1 (Figs. 3c, 3f).

In Experiment 2, we tested a key prediction of the error-based learning mechanism of cue-selective adaptation (Eqn. 5) against an alternative mechanism that involves checking the metric depth estimates derived from each cue against a haptic “ground truth” signal. If an error-based mechanism is responsible for cue-selective adaptation, then it should be observed only during exposure to the stimulus set of Experiment 1 (Fig. 1a; call this an *uncorrelated* set), and not during exposure to a stimulus set where the faulty cue always specifies less depth than the reinforced cue (or more depth; call these *biased* sets). In contrast, if cross-modal metric comparisons allow the system to increase the relative influence of the reinforced cue, then we should observe cue-selective adaptation during exposure to either set.

Biased stimulus sets were comprised of six cue-conflict stimuli where texture depth and stereo depth differed by 10 mm across all objects. The shallower cue always ranged from 20 to 45 mm, the deeper from 30 to 55 mm. We ran two groups of participants in this experiment. For one group (“Adapt+”), the reinforced cue was the deeper of the two cues; for the other group, (“Adapt−”), the reinforced cue was the shallower of the two cues. Each participant performed a haptic-for-texture condition and a haptic-for-stereo condition in separate sessions at least three hours apart. Participants began each session by creating perceptual matches between cue-consistent paraboloids and the six conflict stimuli in the biased set. They then performed grasping movements through five phases: (1) Baseline grasping of the six perceptually matched cue-consistent stimuli; (2) Pre-test grasping of the uncorrelated set from Experiment 1 to estimate cue slopes prior to exposure, with haptic feedback matching the reinforced cue to maintain consistency with Experiment 1; (3) Adaptation grasping of the relevant biased set (10 bins for the Adapt+ group; 5 bins for Adapt−); (4) Post-test grasping of the uncorrelated set from Experiment 1 to estimate cue slopes after exposure, with haptic feedback still remaining consistent with the reinforced cue; and (5) Washout grasping of the perceptually matched cue-consistent stimuli.

Figure 4 depicts the main results of the experiment, with one panel for each feedback condition (haptic-for-texture, haptic-for-stereo) of each group (Adapt+, Adapt−). In the middle of each panel, we present the Baseline-centered average MGAs for each bin of the Adaptation phase (right-hand y-axis, open circles). The dashed red and blue lines spanning the Adaptation phase represent the rendered stereo and texture depths in the biased sets, with the constant 10-mm cue-conflict; one of these cues was consistent with haptic feedback. In Figure 4, the positions of the dashed lines with respect to the average Baseline MGA (zero) reflect the changes in texture and stereo depth from Baseline to Adaptation. Notice that the dashed red line is slightly closer to zero; this is because cue-consistent depths were set closer to the stereo depths than to the texture depths of the cue-conflicts during perceptual matching, consistent with the stronger influence of stereo information on perceived depth.

**Figure 4.**
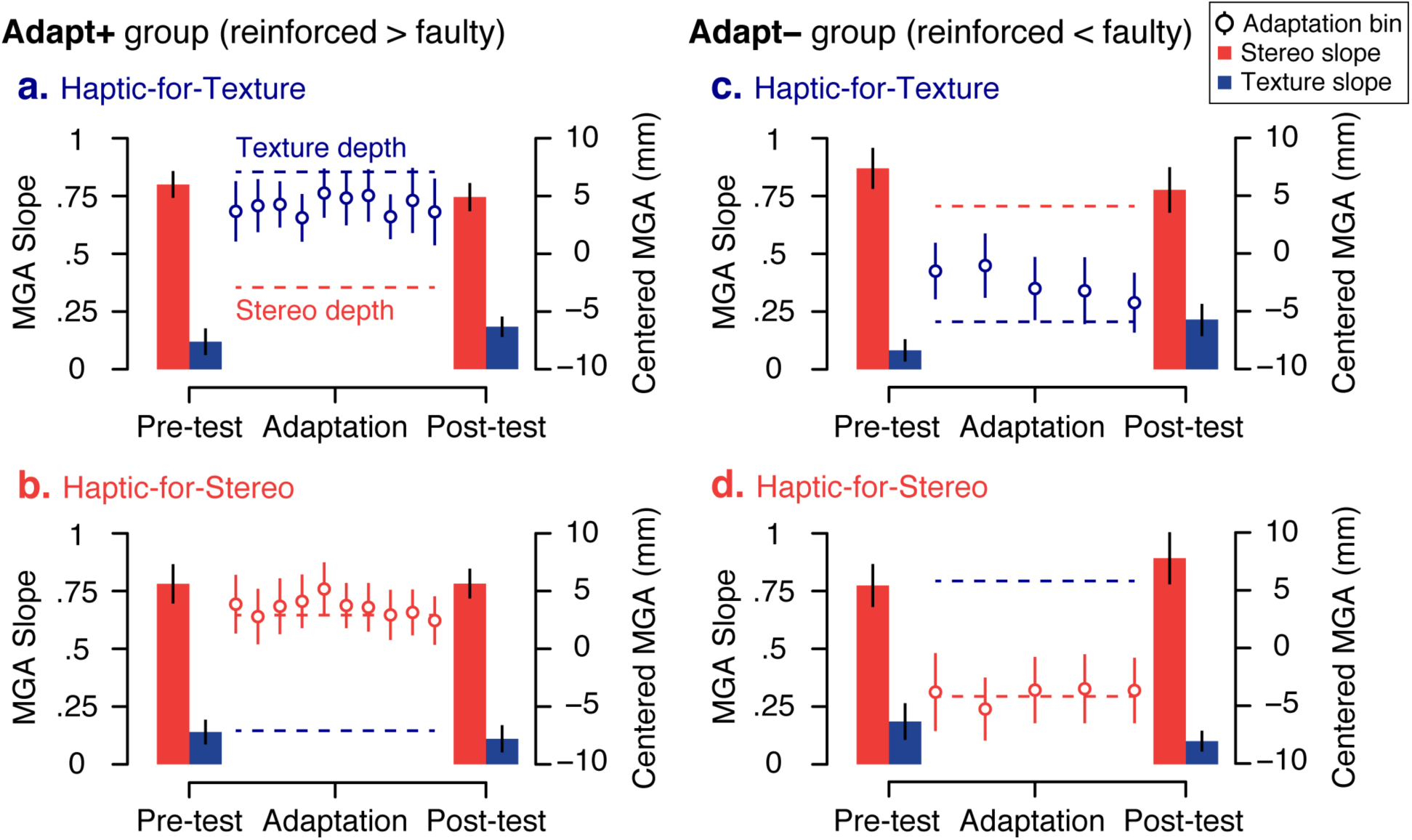
Experiment 2 results. Each panel shows the results of one group-condition pairing. Panels a and b depict the two conditions of the Adapt+ group, while panels c and d show the conditions of the Adapt− group. In the central Adaptation phase, participants grasped objects with a constant cue-conflict: 10-mm separation between texture depth and stereo depth (blue and red dashed lines, respectively). For each bin (6 trials) of the Adaptation phase, we depict the Baseline-centered average maximum grip apertures (MGAs; right y-axis); the symbol color corresponds to the haptically reinforced cue. The length of the Adaptation phase for the Adapt− group was shortened by half based on the rapid adaptation observed for the Adapt+ group. Flanking the main Adaptation phase, the bar graphs indicate the slope of the MGA with respect to stereo and texture information (left y-axis) during the Pre-test and Post-test phases, where we presented the uncorrelated set of stimuli (matrix of Fig. 1).

During the Adaptation phase, MGAs increased (Adapt+: *t*(21) = 4.18, p = 0.00021) or decreased (Adapt−: *t*(17) = 3.21, p = 0.0026) from their Baseline values in order to target the reinforced cue. We were surprised, however, to find that the time course of these data did not reflect the exponential learning curve that is characteristic of adaptation to a constant bias. Even in the very first bin of Adaptation (six trials), grasp planning had already compensated for most or all of the change in the haptic feedback. Originally, we expected to observe a more gradual shifting of the MGAs, as participants in previous grasp adaptation experiments required approximately ten trials to fully adapt in response to similar perturbations (Cesanek & Domini, 2017; Cesanek *et al*., 2019). It is likely that the inclusion of the Pre-test phase between Baseline and Adaptation disrupted the typical time course. In any case, the key result of the Adaptation phase is that MGAs were significantly altered from Baseline, appearing to specifically target the reinforced cue by the end of the phase.

In the Pre-test and Post-test phases, we measured the influences of stereo and texture information during 18 grasps toward the uncorrelated set. Even during these Test trials, haptic feedback remained consistent with the reinforced cue. As in Experiment 1, we estimated a slope parameter for each cue using multiple linear regression (left-hand y-axis, bar graphs). We then performed a mixed-design ANOVA on these slopes with a single between-subjects factor (Group: Adapt+ or Adapt−) and three within-subjects factors (Condition: haptic-for-stereo or haptic-for-texture; Test Phase: Pre-test or Post-test; Cue: stereo or texture). This analysis revealed a highly significant main effect of Cue (*F*(1, 38) = 106.72, p < 0.001), as well as a three-way interaction of Condition ✕ Test Phase ✕ Cue (*F*(1, 38) = 9.41, p = 0.0040).

At first glance, these results appear to suggest that, contrary to our predictions, cue-selective adaptation did in fact take place during exposure to biased set during the Adaptation phase. However, this conclusion overlooks the possibility that these tests captured a gradual accumulation of cue-selective adaptation in the Pre-test and the Post-test. Recall that within each Test phase, participants performed two bins of nine grasps toward the stimuli of the uncorrelated set, with haptic feedback continuing to reinforce the reliable cue in each condition. The exposure during these phases was therefore identical to that in the Adaptation phase of Experiment 1, although considerably briefer. Accordingly, we also evaluated the possibility that cue-selective adaptation occurred *within* the Pre-test and the Post-test, during exposure to the uncorrelated set, and not *across* the central Adaptation phase during exposure the biased sets.

To obtain a finer temporal resolution, we fit multiple linear regressions to measure the influences of stereo and texture information in each of the two bins of Pre-test and Post-test, so we could compare slope changes that occurred *within* the Test phases, from Bin 1 to Bin 2, with those that occurred *across* the Adaptation phase, from Bin 2 of Pre-test to Bin 1 of Post-test (Fig. 5). This is appropriate because Bin 2 of the Pre-test provides the most up-to-date measure of cue influences on grasping prior to any exposure to the biased set. Recall that the error-based update rule in our model predicts no cue-selective adaptation during exposure to the biased set, so the relative influence of the reinforced cue should not be enhanced across the Adaptation phase. On the other hand, we might expect some cue-selective adaptation within the Test phases.

**Figure 5.**
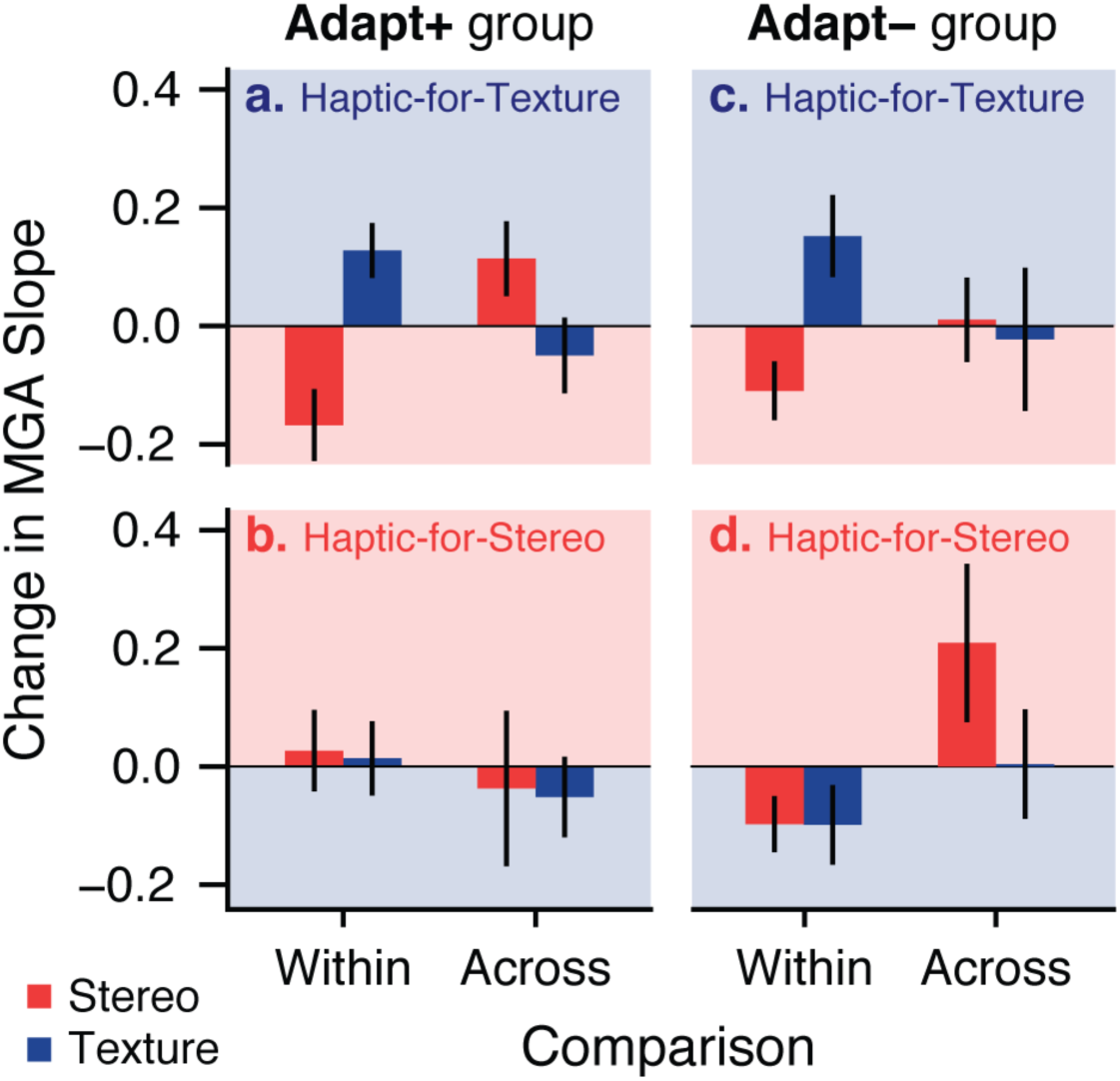
Changes in slope parameters observed *within* the Test phases, as a result of exposure to the uncorrelated set, versus those observed *across* the Adaptation phase, as a result of exposure to the biased sets. The shading of the background indicates the expected direction of slope change for each cue, if cue-selective adaptation took place. For example, in a haptic-for-texture condition, cue-selective adaptation would be marked by an increase in the slope of the MGA with respect to texture information (blue) and/or a decrease in the slope with respect to stereo (red).

First, we used a mixed-design ANOVA as an omnibus test of the slope-change data displayed in Figure 5, with one between-subjects factor, Group (columns of Fig. 5), and two within-subjects factors, Condition (rows of Fig. 5) and Order (x-axis of Fig. 5; changes occurring *within* the Test phases versus those occurring *across* the Adaptation phase). Since cue-selective adaptation is marked by opposing changes in the slopes of the two cues, we have simply taken the difference between the slope changes, change in reinforced minus change in faulty, as our dependent variable. This analysis revealed a significant interaction of Condition ✕ Order (*F*(1,38) = 6.39, p = 0.016), indicating that the within-versus-across difference that we were interested in varied as a function of the feedback condition. Accordingly, we followed up with two specific paired *t*-tests, one for each condition. In the haptic-for-texture condition (Figs. 5a & 5c), we found that cue-selective adaptation was significantly greater within the Test phases than across the Adaptation phase (*t*(39) = 2.92, p = 0.0058). No such difference was found in the haptic-for-stereo condition (p = 0.47)—Figures 5b and 5d reveal mostly negligible slope changes in this condition. An apparent exception can be spotted in Figure 5d (Adapt−, haptic-for-stereo), where it appears that the strength of stereo information did increase considerably across the Adaptation phase. However, on closer inspection we found the Pre-test of this condition to be somewhat anomalous, with unusually low stereo and texture slopes in Bin 2 of Pre-test (0.67 and 0.12, compared to 0.89 and 0.27 in the preceding bin). The low slopes in this bin were accompanied by a very large intercept parameter (39.8 mm, compared to 27.4 mm in the preceding bin), suggesting that participants had adopted a uniformly larger grip aperture and temporarily reduced their normal reliance on depth information. The correction of the abnormally low stereo slopes in Bin 1 of Post-test should not necessarily be taken as evidence of cue-selective adaptation—in fact, a post-hoc test of this subset of the data reveals that the observed increase in stereo slope across Adaptation was not significant (p = 0.075).

Overall, by breaking down our Pre-test and Post-test phases into their constituent bins, we find evidence that the overall cue-selective adaptation effect from Pre-test to Post-test (reported above, see Fig. 4) actually resulted from cumulative exposure to the uncorrelated set in the two Test phases. These data show that the constant perceptual bias introduced during the Adaptation phase was handled by simply increasing or decreasing the grip aperture, rather than changing the relative influences of the cues. Yet during this phase, participants had plenty of exposure to a systematic mismatch between physical depth, felt via haptic feedback, and the depth specified by the faulty visual cue. If a metric, cross-modal comparison of the haptic “ground truth” with each visual cue is capable of reducing the relative influence of the faulty cue in grasp planning, we should observe this as a shift in the regression coefficients from the second bin of Pre-test to the first bin of Post-test, which immediately followed Adaptation. In contrast, we see that exposure to the biased sets produced no cue-selective adaptation. Ultimately, we ended up with additional evidence that exposure to the uncorrelated set, where participants experience conflicting movement errors that are correlated with the faulty cue, is necessary to trigger cue-selective adaptation. These results are consistent with the predictions of our error-based mechanism, and inconsistent with the notion that haptic feedback causes cue-selective adaptation by identifying the inaccurate cue.

## 3. Discussion

Both of the reported experiments reliably induced cue-selective adaptation for 3D targets defined by stereo and texture information, as measured by changes in the slope of the MGA with respect to each cue. This occurred only when the correlation between a given cue and physical reality was altered, and not in response to simple over-or underestimation of depth from that cue, which instead produced a constant shift of the planned grip aperture. As a complement to canonical adaptation, cue-selective adaptation is a powerful mechanism to keep movements accurate when interacting with real 3D objects under variable viewing conditions.

With respect to the neurophysiology of the visuomotor system, perhaps the most notable implication of our findings is that individual depth cues must remain separable in the visual encoding used for motor planning. Cue-selective adaptation would not be possible if available cues were already combined into a single estimate, because movement error signals could not be related back to the input signals received from each cue (Eqn. 5). Thus, we can conclude that texture and stereo signals are separably encoded in the visuomotor mapping.

In apparent contrast to the separable encoding needed to support cue-selective adaptation, a seminal paper from Hillis, Ernst, Banks, and Landy (2002) concluded that “mandatory fusion” of stereo and texture cues occurs in conscious 3D perception. Specifically, their study showed that individual cues to slant cannot be isolated to aid perceptual discriminations between cue-consistent and cue-conflict stimuli. In the remainder of the Discussion, we develop an account of cue-selective adaptation that reconciles the phenomenon of cue-selective adaptation with the idea of mandatory fusion of depth cues in perception. To do so, we will speculate on the relationship between the visual encoding affected by cue-selective adaptation and how perceived 3D shape may emerge from this encoding.

### 3.1. Distributed encoding of depth signals

In the Introduction, we posed the visuomotor mapping as a global function with specific parameters (cue-specific slopes and an intercept) and assumed that these parameters are modified by sensorimotor adaptation. However, some previous research suggests it is more appropriate to model the effects of adaptation using a distributed representation of the visuomotor mapping, where a population of locally-tuned units (“neurons”) encode relevant features of the desired response (*e.g.*, target location, movement velocity; Thoroughman & Shadmehr, 2000; Donchin, Francis, & Shadmehr, 2003; Hwang, Donchin, Smith, & Shadmehr, 2003). In the case of grasping 3D objects defined by multiple depth cues, we are concerned with visual features, namely stereo and texture information. When preparing a grasp, we propose that input signals related to these features establish a pattern of activity across the population of stereo-and texture-sensitive units. For instance, some units might respond maximally when a strong stereo signal is paired with a weak texture signal, while others respond maximally to strong signals from both stereo and texture. With this type of tuning, the units could encode the entire space of possible depth signals, allowing them to produce a response for nearly any cue-specific distortion that might arise. Together, these units and their associated weights specify a particular output at every location in the depth signal space, producing a smooth function akin to, but less constrained than the linear surface depicted in Figure 2.

When planning to grasp a paraboloid object with a large depth-from-stereo and a small depth-from-texture, nodes that are sensitive to this particular combination of visual information may respond preferentially, while nodes sensitive to other combinations are relatively silent. This pattern of activity is then transformed into a planned grip aperture by (1) scaling the input activity of each node by a vector of weight parameters, which determine the influence of each node on the motor output, and (2) pooling the weighted node activities. This approach to computing arbitrary functions is called a local function approximation because the output is constrained to vary smoothly with changes in the input dimensions (stereo and texture information), but unlike Equation 1 there are no explicit global parameters. Instead, the parameters are the node weights, and adjusting the weight of a node based on error feedback will only raise or lower the motor output in a localized region of the input domain. As a result, this format provides a certain amount of selectivity along the feature dimensions encoded by the individual units, that is, it provides the separability of stereo and texture information that is required for cue-selective adaptation.

On this view, cue-selective adaptation is a straightforward result of the same error-correction mechanisms that underlie canonical sensorimotor adaptation, parsimoniously eliminating the *ad hoc* distinction between one form of adaptation that adjusts the intercept and another that adjusts the slope of the visuomotor mapping. In this model, cue-selective adaptation is achieved entirely by incremental, local adjustments of the approximated function. Since the units are selective for specific combinations of depth cue signals, error signals will most strongly affect motor outputs in the region of signal space that encodes the target object, with the corrective effect falling off in more distant regions. As these corrections accumulate, the resulting changes in planned grip apertures can be captured in the parameters estimated by multiple linear regression. Intercept shifts occur when node weights all change in the same direction, as when experiencing constant errors regardless of the available depth information. On the other hand, slope changes occur when node weights change in opposite directions in different regions of the input space, as when large values of an unreliable depth cue cause positive errors and small values cause negative errors.

### 3.2. Cue-selective adaptation and 3D perception

In this study, we did not test whether the observed cue-selective adaptation is actually the result of a deeper change in 3D visual perception. However, a previous study by Ernst *et al*. (2000) suggests that the same kind of visuomotor training does produce changes in perception of 3D slant. In that study, participants touched a variety of slanted surfaces in which either stereo or texture information was consistent with the physical surface slant; the other (unreliable) cue varied dramatically around the physical slant (±30°). After exposure, participants’ slant judgments were less influenced by the unreliable cue. The authors, seeking to justify their “small” observed changes in perception, point out that their unreliable cue remained somewhat correlated with physical slant (*r* = 0.59). However, this may be a misleading view of the experimental conditions, as their stimuli would have actually produced a strong correlation of the unreliable cue with movement error signals, which we have suggested as the driver of cue-selective adaptation according to the learning rule of Equation 5 (note that this update rule is perfectly suited for tuning node weights in the distributed representation discussed above). Given the highly similar patterns of error feedback in the present study and the study of Ernst *et al*. (2000), an intriguing possibility is that the process of cue-selective adaptation actually produces changes in visual perception, and not just changes in the motor output.

Though this is a speculative proposal, consistency with empirical results indicates that it is not unreasonable to believe that perceived depth is shaped by the process of tuning weights on a population of visual units, as described in the previous section. Specifically, we proposed that the units in this population are responsive to particular *combinations* of signals from earlier stereo processing and texture processing. To explain the results of Ernst *et al*. (2000), we propose that the adjustment of weights on these units does not merely modify the motor output, like the planned grip aperture, but actually determines the resulting perceptual interpretation of the present assortment of depth information. Intriguingly, this view also fits with the mandatory fusion of depth cues in perceptual judgments, as demonstrated by Hillis *et al*. (2002). In creating a particular pattern of activity across the full population of units, available depth cues have already been combined—although the pattern of activity does indicate a specific combination of cue signals, which provides separability, the component signals can no longer be individually accessed. Therefore, if perceived 3D shape is determined by the learned “weights” on the presently activated units, which connect them to downstream systems for action planning, then when two distinct combinations of depth signals activate units with similar weights, the resulting percepts will not be discriminable, even if the individual cues might be discriminable when presented in isolation.

### 3.3. The teaching signal: Haptic information versus movement errors

As a final note, it is also worth pointing out that the earlier connection with the study by Ernst *et al*. (2000) helps to illuminate a subtle but important difference between movement errors and haptic estimates of 3D shape as drivers of cue-selective adaptation. Haptic estimates of physical shape are often suggested as a ground truth against which to calibrate visual processing of depth information, following Berkeley’s maxim that “touch teaches vision”. The gist of the claim is that the relative reliabilities of depth cues can be estimated by tracking correlations between haptic estimates of depth and visual estimates of depth obtained independently from each cue. In practice it would be extremely difficult to robustly estimate the relative reliabilities of two noisy signals, stereo and texture, based on their correlations with a third, even noisier signal, haptics. However, many authors have assumed that relative reliabilities can be accurately estimated by comparison with haptic information and subsequently leveraged to combine the individual depth estimates from each cue in statistically optimal fashion (Ernst & Banks, 2002; Hillis, Watt, Landy, & Banks, 2004). Perhaps a bigger issue with this mechanism for cue combination is that it depends on yet another questionable assumption, that individual cues provide unbiased metric estimates of physical depth. There is a large body of psychophysical evidence that directly conflicts with this assumption (Todd, 2004).

In contrast, the learning rule we have suggested for adjusting the interpretation of different combinations of depth cue signals (Eqn. 5) could be driven by the same error signals that have been implicated in sensorimotor adaptation and does not require any controversial assumptions about the availability of unbiased estimates. Rather, it represents a way in which naturally occurring fluctuations in the reliability of individual depth cues could be handled by the sensorimotor system, through iterative online adjustments that gradually reduce the influence of signals that are positively correlated with error. The end result is an interpretation of depth information from multiple cues, each of which may be expressed in arbitrary units, that reflects the distinctions that are most important for producing accurate actions. This may be viewed as a less intelligent mechanism of cue combination, as it does away with the idea that the visual system recovers accurate depth estimates from each individual cue. However, this intelligence, inherited from the naïve realist view that perception is firmly moored in the metrics of physical reality, is replaced by a powerful short-term flexibility that allows perception to effectively drive action across changing environmental conditions.

## 4. Methods

### 4.1. Participants

Sixty-five participants were recruited for Experiments 1 (N = 25) and 2 (N=40; 22 in Adapt+, 18 in Adapt−). Participants were between 18 and 35 years old and right-handed, with normal or corrected-to-normal vision. They were either granted course credit or paid $8/hour as compensation. Informed consent was obtained from all participants prior to any participation, in accordance with protocol approved by the Brown University Institutional Review Board and performed in accordance with the ethical standards set forth in the Declaration of Helsinki.

### 4.2. Apparatus

Figure S1 presents a few photographs of the lab setup. Participants were seated in a height-adjustable chair so that the chin rested comfortably in a chinrest. Movements of the right hand were tracked using an Optotrak Certus motion-capture system. Small, lightweight posts containing three infrared emitting diodes were attached to the fingernails of the index finger and thumb, and the system was calibrated prior to the experiment to track the extreme tips of the distal phalanxes of each finger. This motion-capture system was coupled to a tabletop virtual reality environment: participants looked into a half-silvered mirror slanted at 45° relative to the sagittal body midline, which reflected the image displayed on a 19” CRT monitor placed directly to the left of the mirror at the correct distance to provide consistent accommodative and vergence information.

Participants viewed stereoscopic renderings of 3D paraboloid objects in this tabletop virtual reality setup, where stereo and texture information were controlled independently using a 3D graphics technique known as backprojection. The paraboloids were rendered with their tips at a viewing distance of 40 cm at eye level. This arrangement made the rendered 3D objects appear to be floating in space beyond the mirror. The bases of the paraboloids always subtended 6.5° of visual angle. Stereoscopic presentation was achieved with a frame interlacing technique in conjunction with liquid-crystal goggles synchronized to the frame rate. Stereoscopic visual feedback of the thumb only was provided throughout the experiment, to help participants keep track of their hand position. We presented the thumb only in order to prevent visual comparison of the stereo-rendered grip aperture with the stereo-specified depth of the object, which might have unintentionally reinforced stereo information in our haptic-for-texture conditions.

To provide haptic feedback, a custom-built motorized apparatus was placed in the workspace. This apparatus consisted of a stepper motor with its shaft extended by a long screw. On the end of this screw, we attached a round metal nub to simulate the rounded tip of the paraboloid objects—perfect alignment between the physical and rendered paraboloid tips was established during the calibration phase at the start of each session. To simulate the flat, round rear end of the paraboloids, we threaded a metal washer (approximately 6 cm in diameter, equal to the average base diameter of the rendered objects) onto the screw. As the stepper motor spun, the washer would travel back and forth along the length of the screw, anchored by a heavy wooden piece on the left side to ensure that one full rotation of the stepper motor would move the washer by one thread pitch (*i.e.*, the distance between threads). On every trial, the resulting depth of the physical object was double-checked using additional Optotrak markers mounted on the physical apparatus and corrected if necessary.

### 4.3. Procedure

Both experiments began with a Matching task, where participants matched through a psychometric procedure the perceived depth of the cue-conflict paraboloids with cue-consistent objects, where stereo and texture specified the same depth. In Experiment 1, the target stereo-texture conflict stimuli were the six off-diagonal objects in our uncorrelated set (see Fig. 1 for more details); the three cue-consistent objects of the uncorrelated set did not require matching because they were already composed of consistent cues. In Experiment 2, the target stimuli were six cue-conflict objects where stereo depth and texture depth differed by a constant conflict of 10 mm (see Section 2.2 for more details). On each Matching trial, we allowed participants to switch freely between the fixed cue-conflict stimulus and an adjustable cue-consistent stimulus, using keypresses to make incremental changes to the depth of the cue-consistent stimulus until it appeared to match the depth of the cue-conflict stimulus. To prevent the use of motion information, we displayed a blank screen with a small fixation dot for an inter-stimulus interval of 750 ms whenever the stimulus was changed. Participants performed two repetitions on each of the six fixed cue-conflicts in each Experiment, for a total of 12 Matching trials.

The resulting sets of stimuli (six pairs of matched cue-conflict and cue-consistent paraboloids) were then presented as stimuli in the Baseline, Adaptation, and Washout phases of the Grasping task. During the Grasping task, participants used a precision grip to grasp the paraboloid objects from front to back. Trials were presented in a pseudo-random “binned” trial order, where each of the target objects in a given phase of the experiment was presented once before any one was presented again; as a result, each bin contains one presentation of each target object. Since there were nine target objects in the uncorrelated set, each bin of Experiment 1 contained nine trials. Since there were six target objects in the biased sets, each bin of Experiment 2 contained six trials, except in the Pre-test and Post-test phases where we presented the uncorrelated set. On each trial, participants were shown the target object for 500 ms, then heard the “go” signal, and reached to grasp the target as shown in Fig. S1. There was no explicit time limit on these grasps, but the total elapsed time from movement onset to object contact rarely exceeded 1.5 seconds.

Following the Matching task, the Grasping procedure of Experiment 1 was as follows. In the Baseline phase, participants grasped their personalized set of nine cue-consistent paraboloids, perceptually matched to the nine objects of the uncorrelated set, for three bins of trials. Participants then proceeded immediately into the Adaptation phase, where the cue-consistent paraboloids were suddenly replaced by the perceptually matched cue-conflict paraboloids, and the depth of the physical object providing haptic feedback matched the depth of whichever cue was being reinforced during that session (haptic-for-stereo or haptic-for-texture). Following eleven bins of exposure to the uncorrelated set, Experiment 1 concluded with a two-bin Washout phase, identical to Baseline.

The procedure of Experiment 2 was designed to be similar to Experiment 1, but with presentations of the biased set, with its constant cue-conflict of 10 mm, rather than the uncorrelated set of Experiment 1, with its variable positive and negative cue-conflicts. As in Experiment 1, participants began with a Baseline phase, grasping their personalized set of six cue-consistent paraboloids, which were perceptually matched to the six objects of the biased set, for three bins of trials. Instead of proceeding directly into the Adaptation phase, where they would interact with the cue-conflict objects of the biased set, they first completed a Pre-test phase consisting of two bins of trials where we presented the uncorrelated set. Next, in the Adaptation phase, we presented the six objects of the biased set for 10 bins of trials when testing the Adapt+ group, but only for 5 bins of trials when testing the Adapt− group. We shortened the Adaptation phase for Adapt− because this version of the experiment was run after the Adapt+ group, where we already observed rapid convergence on the reinforced cue—longer adaptation periods were clearly not necessary to eliminate movement errors. Following Adaptation, participants completed a two-bin Post-test, identical to the Pre-test, and concluded with a two-bin Washout phase, identical to Baseline.

### 4.4. Analysis

Raw motion-capture position data was processed and analyzed offline using custom software. Missing frames due to marker dropout were linearly interpolated, and the 85-Hz raw data was smoothed with a 20-Hz low-pass filter. The time series data from each trial was cropped by defining the start frame as the final frame where the thumb was more than 25 cm from its contact location on the tip of the object, and the end frame as the first frame where (a) the thumb came within 1 cm of its contact location or (b) the index finger entered into a 3 cm wide by 3 cm high bounding box, extending 10 cm in depth (well beyond the rear edge of the deepest object). The grip aperture profile was computed for each trial by taking the vector distance between the index finger and thumb locations on each frame. The maximum grip aperture (MGA), a widely used kinematic measure of grasp planning (Jeannerod, 1981), was extracted from this time series.

Two criteria were used for trial exclusion: the proportion of missing frames due to marker dropout exceeded 90% or fewer than 5 frames were not missing. In Experiment 1, neither of these criteria were met for any of the movements, so no trials were excluded from analysis. In Experiment 2, 72 out of a total 9480 trials were excluded by these criteria (~0.7%).

The factorial design of the uncorrelated set of stimuli allowed us to measure the relative influence of stereo and texture information in the Grasping task by estimating coefficients (slopes) for each cue via multiple linear regression according to Equation 1, with the MGA as the response variable. A regression was computed for each bin of nine trials within the Adaptation phase of Experiment 1, and in the Pre-test and Post-test phases of Experiment 2. In the Baseline phase of Experiment 1, we computed the slope of the MGA with respect to the perceptually matched cue-consistent depths using simple linear regression in each bin.

## Acknowledgments

This work was supported by a National Science Foundation grant to F. D. (#BCS 1827550).

---

**Figure S1.**
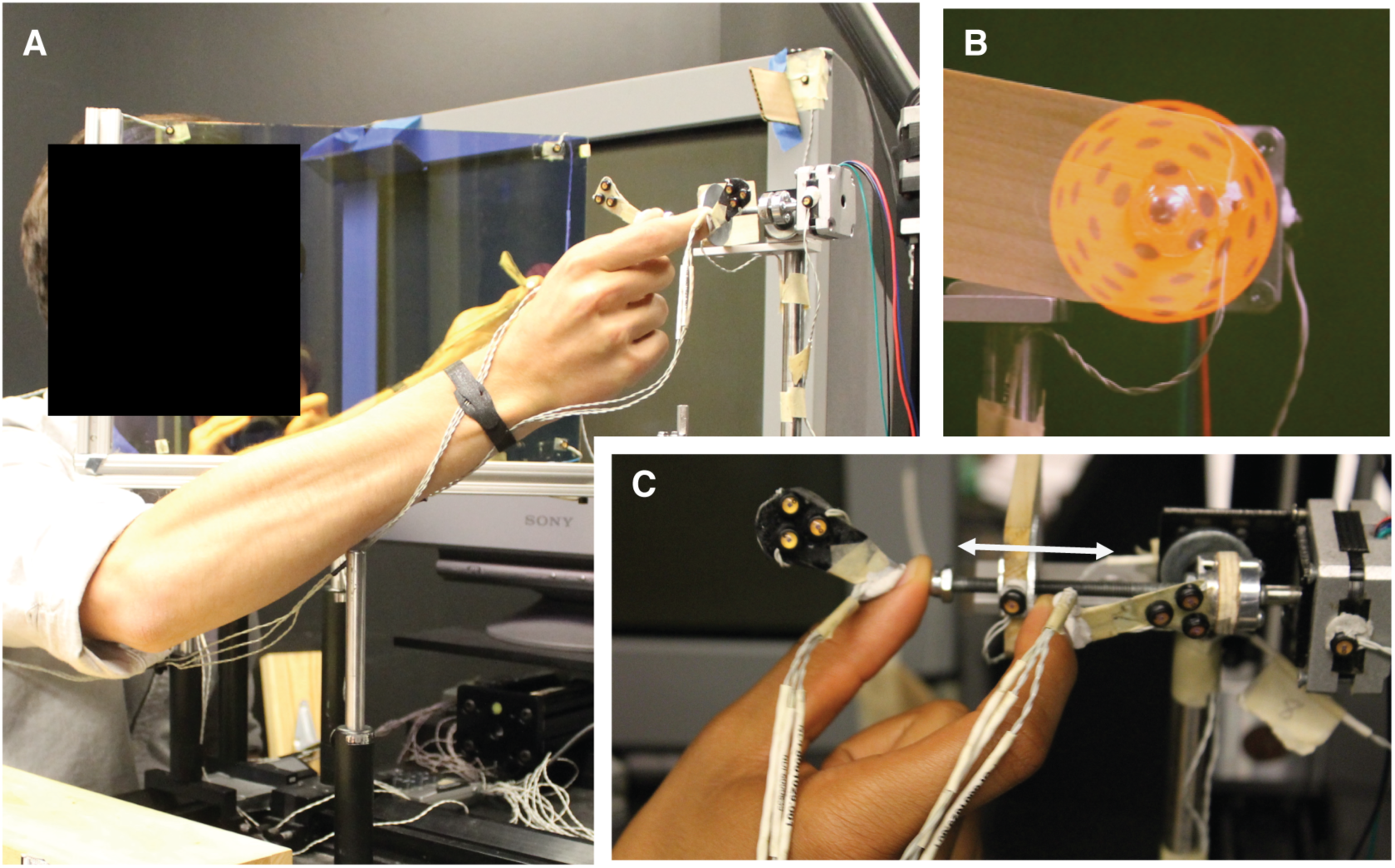
Photographs of the tabletop virtual reality setup. (A) The observer (face hidden for pre-print) looks into a slanted mirror while wearing stereoscopic glasses, seeing a compelling 3D object on the far side of the mirror, aligned with an automated physical apparatus in the workspace that provides haptic feedback of objects with different depths. During the experiment, the room is completely dark and a back panel is placed on the mirror to prevent any visual stimulation other than the rendered 3D object. The participant reaches with the right hand to grasp the rendered object so the thumb lands on the tip and the index finger lands on the base. Optotrak infrared-emitting diodes attached to the fingernails provide precise location information about the fingertips, allowing us to compute the in-flight grip aperture. (B) Frontal view of a rendered paraboloid. This is a cyclopean view, rather than stereoscopic, for visualization purposes. The tip of the paraboloid is perfectly centered on a small rounded nub to provide haptic feedback of the tip. (C) Side view of the physical apparatus for providing haptic feedback. A stepper motor spins a screw in order to slide a large round washer back and forth along the screw. This allowed us to create a physical object of any depth on each trial. The thumb landed on the rounded nub aligned with the tip of the paraboloid, while the index finger pinched down on the rear surface, which could be aligned with either the stereo-or texture-specified depth when grasping a cue-conflict paraboloid.

## References

Cesanek, E. & Domini, F. (2017). Error correction and spatial generalization in human grasp control. Neuropsychologia, 106, 112–122.

Cesanek, E., Taylor, J. A., & Domini, F. (2019). Sensorimotor adaptation compensates for distortions of 3D shape information. bioRxiv. doi: 10.1101/540187.

Cheng, S. & Sabes, P. N. (2006). Modeling sensorimotor learning with linear dynamical systems. Neural Computation, 18(4), 760–793.

Donchin, O., Francis, J. T., & Shadmehr, R. (2003). Quantifying generalization from trial-by-trial behavior of adaptive systems that learn with basis functions: Theory and experiments in human motor control. Journal of Neuroscience, 23(27), 9032–9045.

Ernst, M. O. & Banks, M. S. (2002). Humans integrate visual and haptic information in a statistically optimal fashion. Nature, 415(6870), 429–433.

Ernst, M. O., Banks, M. S., & Bülthoff, H. H. (2000). Touch can change visual slant perception. Nature Neuroscience, 3(1), 69–73.

Held, R. & Hein, A. V. (1958). Adaptation of disarranged hand-eye coordination contingent upon re-afferent stimulation. Perceptual and Motor Skills, 8(3), 87–90.

von Helmholtz, H. E. F. (1962). Treatise on physiological optics. (J. P. C. Southall, Ed. and Trans.). New York: Dover. (Original work completed in 1867, published in 1909).

Hillis, J. M., Ernst, M. O., Banks, M. S., & Landy, M. S. (2002). Combining sensory information: mandatory fusion within, but not between, senses. Science, 298(5598), 1627–1630.

Hillis, J. M., Watt, S. J., Landy, M. S., & Banks, M. S. (2004). Slant from texture and disparity cues: Optimal cue combination. Journal of Vision, 4(12), 967–992.

Hwang, E. J., Donchin, O., Smith, M. A., & Shadmehr, R. (2003). A gain-field encoding of limb position and velocity in the internal model of arm dynamics. PLoS Biology, 1(2), 209–220.

Jeannerod, M. (1981). Intersegmental coordination during reaching at natural visual objects. In: J. Long and A. Baddeley (Eds.), Attention and Performance IX (pp. 153–168). Hillsdale, NJ: Lawrence Erlbaum Associates.

Johnston, E. B. (1991). Systematic distortions of shape from stereopsis. Vision Research, 31(7-8), 1351–1360.

Krakauer, J. W., Ghez, C., & Ghilardi, M. F. (2005). Adaptation to visuomotor transformations: Consolidation, interference, and forgetting. Journal of Neuroscience, 25(2), 473–478.

Krakauer, J. W., Pine, Z. M., Ghilardi, M. F., & Ghez, C. (2000). Learning of visuomotor transformations for vectorial planning of reaching trajectories. Journal of Neuroscience, 20(23), 8916–8924.

Pouget, A. & Snyder, L. H. (2000). Computational approaches to sensorimotor transformations. Nature Neuroscience, 3(11s), 1192–1198.

Redding, G. M. & Wallace, B. (1997). Adaptive spatial alignment. Mahwah, NJ: Lawrence Erlbaum Associates.

Rosenholtz, R. & Malik, J. (1997). Surface orientation from texture: isotropy or homogeneity (or both)?. Vision Research, 37(16), 2283–2294.

Säfström, D. & Edin, B. B. (2005). Short-term plasticity of the visuomotor map during grasping movements in humans. Learning & Memory, 12(1), 67–74.

Säfström, D. & Edin, B. B. (2008). Prediction of object contact during grasping. Experimental Brain Research, 190(3), 265–277.

Smeets, J. B. & Brenner, E. (1999). A new view on grasping. Motor Control, 3(3), 237–271.

Taylor, J. A., Krakauer, J. W., & Ivry, R. B. (2014). Explicit and implicit contributions to learning in a sensorimotor adaptation task. Journal of Neuroscience, 34(8), 3023–3032.

Thoroughman, K. A. & Shadmehr, R. (2000). Learning of action through adaptive combination of motor primitives. Nature, 407(6805), 742–747.

Todd, J. T. (2004). The visual perception of 3D shape. Trends in Cognitive Sciences, 8(3), 115–121.

Welch, R. B. (1969). Adaptation to prism-displaced vision: The importance of target-pointing. Perception & Psychophysics, 5(5), 305–309.

Welch, R. B. (1978). Perceptual modification: Adapting to altered sensory environments. New York, NY: Academic Press, Inc.

